# Gene flow between divergent cereal‐ and grass-specific lineages of the rice blast fungus *Magnaporthe oryzae*

**DOI:** 10.1101/161513

**Authors:** Pierre Gladieux, Bradford Condon, Sebastien Ravel, Darren Soanes, Joao Leodato Nunes Maciel, Antonio Nhani, Li Chen, Ryohei Terauchi, Marc-Henri Lebrun, Didier Tharreau, Thomas Mitchell, Kerry F. Pedley, Barbara Valent, Nicholas J. Talbot, Mark Farman, Elisabeth Fournier

**Author notes:** These authors contributed equally to the work (Pierre Gladieux, Bradford Condon). Corresponding author: Elisabeth Fournier.

## Abstract

Delineating species and epidemic lineages in fungal plant pathogens is critical to our understanding of disease emergence and the structure of fungal biodiversity, and also informs international regulatory decisions. *Pyricularia oryzae (syn. Magnaporthe oryzae)* is a multi-host pathogen that infects multiple grasses and cereals, is responsible for the most damaging rice disease (rice blast), and of growing concern due to the recent introduction of wheat blast to Bangladesh from South America. However, the genetic structure and evolutionary history of *M. oryzae*, including the possible existence of cryptic phylogenetic species, remain poorly defined. Here, we use whole-genome sequence information for *76 M. oryzae* isolates sampled from 12 grass and cereal genera to infer the population structure of *M. oryzae*, and to reassess the species status of wheat-infecting populations of the fungus. Species recognition based on genealogical concordance, using published data or extracting previously-used loci from genome assemblies, failed to confirm a prior assignment of wheat blast isolates to a new species (*Pyricularia graminis tritici*). Inference of population subdivisions revealed multiple divergent lineages within *M. oryzae*, each preferentially associated with one host genus, suggesting incipient speciation following host shift or host range expansion. Analyses of gene flow, taking into account the possibility of incomplete lineage sorting, revealed that genetic exchanges have contributed to the makeup of multiple lineages within *M. oryzae*. These findings provide greater understanding of the eco-evolutionary factors that underlie the diversification of *M. oryzae* and highlight the practicality of genomic data for epidemiological surveillance in this important multi-host pathogen.

**Importance:** Infection of novel hosts is a major route for disease emergence by pathogenic micro-organisms. Understanding the evolutionary history of multi-host pathogens is therefore important to better predict the likely spread and emergence of new diseases. *Magnaporthe oryzae* is a multi-host fungus that causes serious cereal diseases, including the devastating rice blast disease, and wheat blast, a cause of growing concern due to its recent spread from South America to Asia. Using whole genome analysis of 76 fungal strains from different hosts, we have documented the divergence of *M. oryzae* into numerous lineages, each infecting a limited number of host species. Our analyses provide evidence that inter-lineage gene flow has contributed to the genetic makeup of multiple *M. oryzae* lineages within the same species. Plant health surveillance is therefore warranted to safeguard against disease emergence in regions where multiple lineages of the fungus are in contact with one another.

## Introduction

Investigating population genetic structure in relation to life history traits such as reproductive mode, host range or drug resistance is particularly relevant in pathogens [1, 2]. Knowledge of species, lineages, populations, levels of genetic variability and reproductive mode is essential to answer questions common to all infectious diseases, such as the tempo, origin, proximate (i.e. molecular) and ultimate (eco-evolutionary) causes of disease emergence and spread [3]. Multilocus molecular typing schemes have shown that cryptic species and lineages within species are often more numerous than estimated from phenotypic data alone. Genomic approaches are emerging as a new gold standard for detecting cryptic structure or speciation with increased resolution, allowing fine-grained epidemiological surveillance and science-based regulatory decisions. The added benefit of whole genomes approaches includes identifying the genetic basis of life history traits, and better understanding of both the genomic properties that influence the process of speciation and the signatures of (potentially incomplete) speciation that are observable in patterns of genomic variability [4, 5].

Many plant pathogenic ascomycete fungi are host-specific, and some of their life history traits have been shown to be conducive to the emergence of novel pathogen species adapted to new hosts [6, 7]. Investigating population structure within multi-host ascomycetes thus offers a unique opportunity to identify the genomic features associated with recent host-range expansions or host-shifts. In this study, our model is Magnaporthe oryzae (*synonym of Pyricularia oryzae*) [6-8], a fungal ascomycete causing blast disease on a variety of grass hosts. *Magnaporthe oryzae* is well studied as the causal agent of the most important disease of rice (*Oryza sativa*), but it also causes blast disease on more than 50 cultivated and wild monocot plant species [9]. This includes other cereal crops such as wheat (*Triticum aestivum*), barley (*Hordeum vulgare*), finger millet (*Eleusine coracana*), and foxtail millet (*Setaria italica, S. viridis*), as well as wild and cultivated grass hosts including goosegrass (*Eleusine indica*), annual ryegrass (*Lolium multiflorum*), perennial ryegrass (*L. perenne*), tall fescue (*Festuca arundinacea*) and St. Augustine grass (*Stenotaphrum secundatum*) [10]. Previous studies based on multilocus sequence typing showed that *M. oryzae* is subdivided into multiple clades, each found on only a limited number of host species, with pathogenicity testing revealing host-specificity as a plausible driver of genetic divergence [11, 12]. More recently, comparative genomics of eight isolates infecting wheat, goosegrass, rice, foxtail millet, finger millet and barley revealed deep subdivision of *M. oryzae* into three groups infecting finger millet or wheat, foxtail millet, and rice or barley [13, 14]. Subsequent analysis of genomic data from nine wheat-infecting isolates, two ryegrass infecting isolates, and one weeping lovegrass-infecting isolate subdivided lineages infecting wheat only on the one hand and wheat or ryegrass on the other hand, and revealed an additional lineage associated with the weeping lovegrass strain [15]. Together, these studies suggest a history of host-range expansion or host-shifts and limited gene flow between lineages within *M. oryzae.*

*Magnaporthe oryzae* isolates causing wheat blast represent a growing concern in terms of food security. This seed-borne pathogen can spread around the world through movement of seed or grain. Therefore, understanding the evolutionary origin and structure of populations causing wheat blast is a top priority for researchers studying disease emergence and for regulatory agencies. Wheat blast was first discovered in southern Brazil in 1985 [16] and the disease subsequently spread to the neighboring countries of Argentina, Bolivia and Paraguay [17-19] where it represents a considerable impediment to wheat production [20, 21]. Until recently, wheat blast had not been reported outside South America. In 2011, a single instance of infected wheat was discovered in the U.S., but analysis of the isolate responsible revealed that it was genetically similar to a local isolate from annual ryegrass and, therefore, unlikely to be an exotic introduction from South America [22]. More recently, in 2016, wheat blast was detected in Bangladesh [23]. Unlike the U.S. isolate, strains from this outbreak resembled South American wheat blast isolates rather than ryegrass-derived strains [15, 23], thereby confirming the spread of wheat blast from South America to Bangladesh.

It has recently been proposed that a subgroup of the wheat-infecting isolates, together with some strains pathogenic on *Eleusine* spp. and other *Poaceae* hosts, belongs to a new phylogenetic species, *Pyricularia graminis-tritici* (*Pgt*) that is well separated from other wheat‐ and ryegrass-infecting isolates, as well as pathogens of other grasses [24]. However, this proposed split was based on bootstrap support in a genealogy inferred from multilocus sequence concatenation, and Genealogical Concordance for Phylogenetic Species Recognition was not applied (GCPSR; [25, 26]). The observed lineage divergence appeared to be mostly driven by genetic divergence at one of 10 sequenced loci, raising questions on the phylogenetic support of this species.

The present study was designed to re-assess the hypothesis that *Pgt* constitutes a cryptic species within *M. oryzae* and, more generally, to infer population structure in relation to host of origin in this important plant pathogen. Using whole-genome sequences for 81 *Magnaporthe* isolates (*76 M. oryzae* from 12 host plant genera, four *M. grisea* from crabgrass [*Digitaria spp.*], and one *M. pennisetigena from Pennisetum* sp.) we addressed the following questions: do *M. oryzae* isolates form distinct host-specific lineages; and is there evidence for relatively long-term reproductive isolation between lineages (i.e. cryptic species) within *M. oryzae*? Our analyses of population subdivision and species identification revealed multiple divergent lineages within *M. oryzae*, each preferentially associated with one host plant genus, but refuted the existence of a novel cryptic phylogenetic species named *P. graminis-tritici*. In addition, analyses of gene flow revealed that genetic exchanges have contributed to the makeup of the multiple lineages within *M. oryzae.*

## Results

### Re-assessing the validity of the proposed *P. graminis-tritici* species by analyzing the original published data according to Phylogenetic Species Recognition by Genealogical Concordance (GCPSR)

To test the previous delineation of a subgroup of wheat-infecting isolates as a new phylogenetic species, we re-analyzed the Castroagudin et al. dataset [24], which mostly included sequences from Brazilian isolates. However, instead of using bootstrap support in a total evidence genealogy inferred from concatenated sequences for species delineation, we applied the GCPSR test [25, 26]. This test identifies a group as an independent evolutionary lineage (i.e. phylogenetic species) if it satisfies two conditions: (1) Genealogical concordance: the group is present in the majority of the single-locus genealogies, (2) Genealogical nondiscordance: the group is well-supported in at least one single-locus genealogy and is not contradicted in any other genealogy at the same level of support [25]. Visual inspection of the topologies and supports in each single-locus tree revealed that GCPSR condition (1) was not satisfied since isolates previously identified as belonging to the phylogenetic species *Pgt* grouped together in only one maximum likelihood gene genealogy – the one produced using the *MPG1* locus (Figure S1A). The *Pgt* separation was not supported by any of the nine other single-locus genealogies (Figures S1B-S1J).

Next, we used the multilocus data as input to the program ASTRAL with the goal of inferring a species tree that takes into account possible discrepancies among individual gene genealogies [27-29]. The ASTRAL tree failed to provide strong support for the branch holding the isolates previously identified as *Pgt* (Figure S2). Thus, analysis of the Castroagudin et al. data according to GCPSR standards failed to support the existence of the newly described *Pgt* species.

### Inferring population subdivision within *M. oryzae* using whole genome data

We sought to test whether a phylogenomic study could provide better insight into the possibility of speciation within *M. oryzae*. To this end, whole genome sequence data were acquired for a comprehensive collection of 76 *M. oryzae* isolates from 12 host genera, four *M. grisea* isolates from *Digitaria* spp. and one *M. pennisetigena* isolate from *Pennisetum* (Table 1). The analysis included sequence data for strains collected on rice (*Oryza sativa*), finger millet and goosegrass (*Eleusine spp.*), wheat (*Triticum spp.*), tall fescue (*Festuca arundinaceum*), annual and perennial ryegrasses (*Lolium multiflorum and L. perenne*, respectively), and barley (*Hordeum vulgare*). Representatives of previously unstudied host-specialized populations from foxtails (*Setaria sp.*), St. Augustine grass (*Stenotaphrum secundatum*), weeping lovegrass (*Eragrostis curvula*), signalgrass (*Brachiaria sp.*), cheatgrass (*Bromus tectorum*) and oat (*Avena sativa*) were also included. SNPs identified in aligned sequences of 2,682 orthologous single copy genes identified in all *M. oryzae* genomes (in total ~6.6 Mb of sequence data), and from whole-genome SNPs identified from pairwise blast alignments of repeat-masked genomes (average ~36 Mb aligned sequence).

**Table 1.**
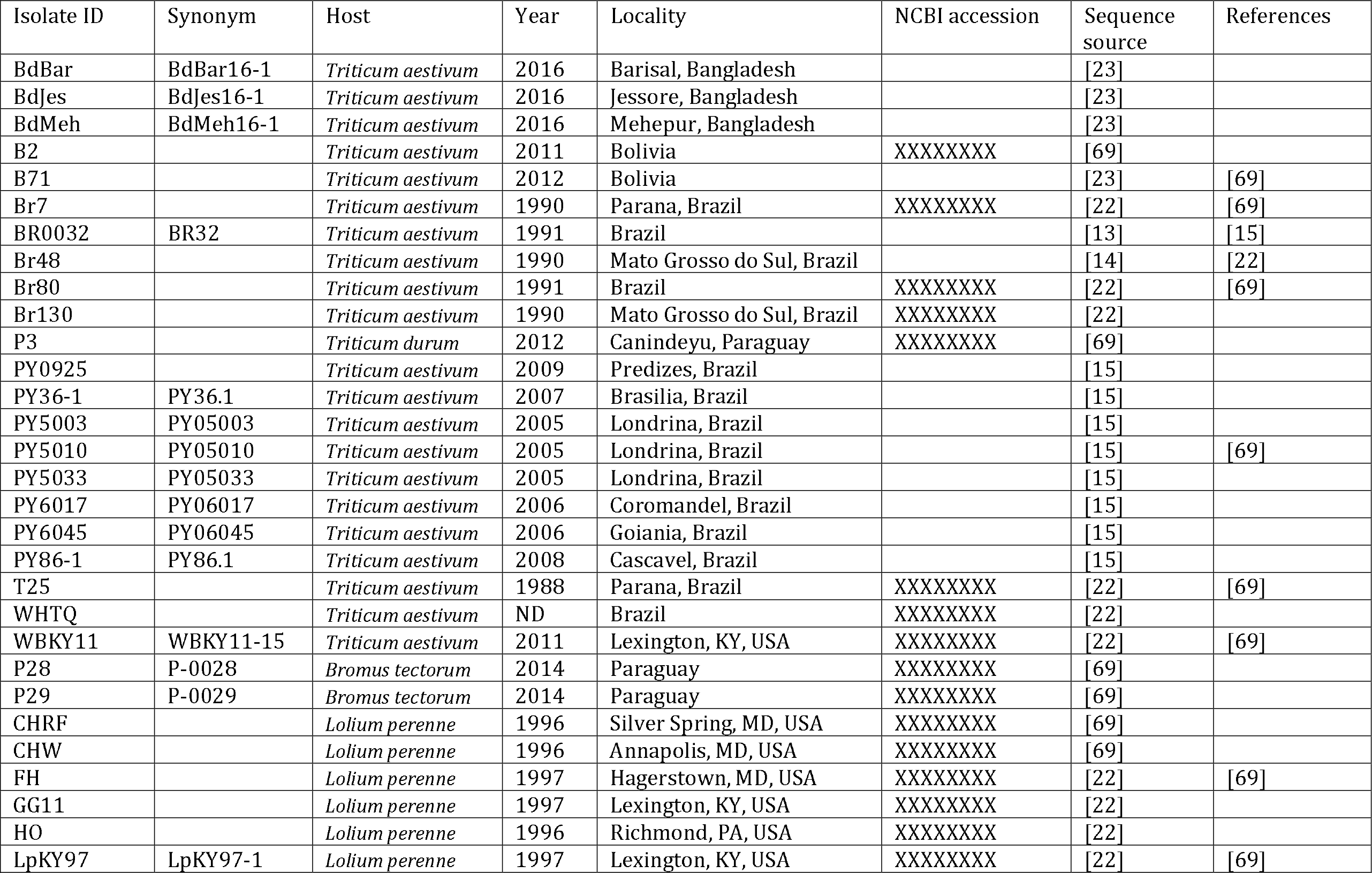
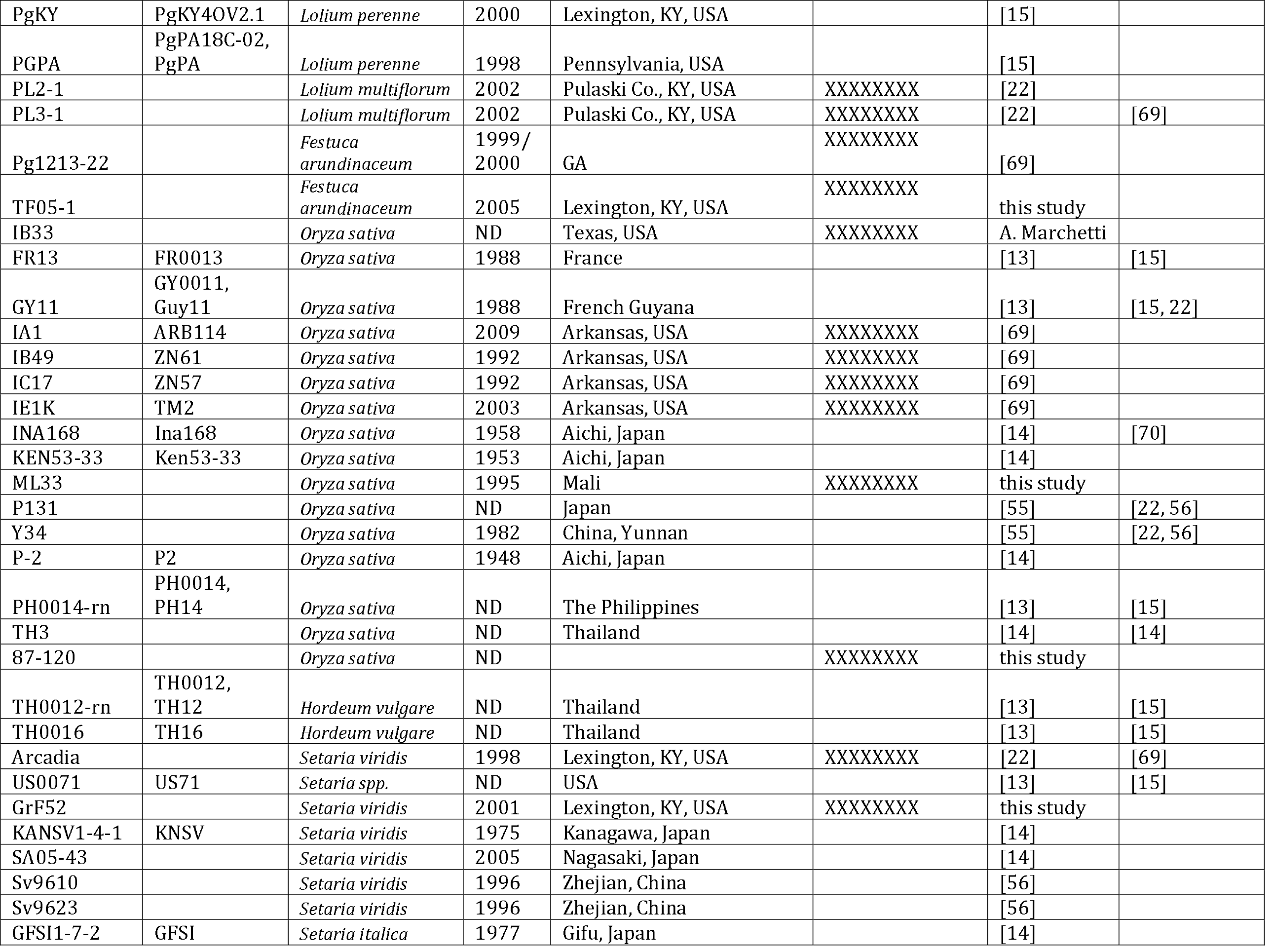
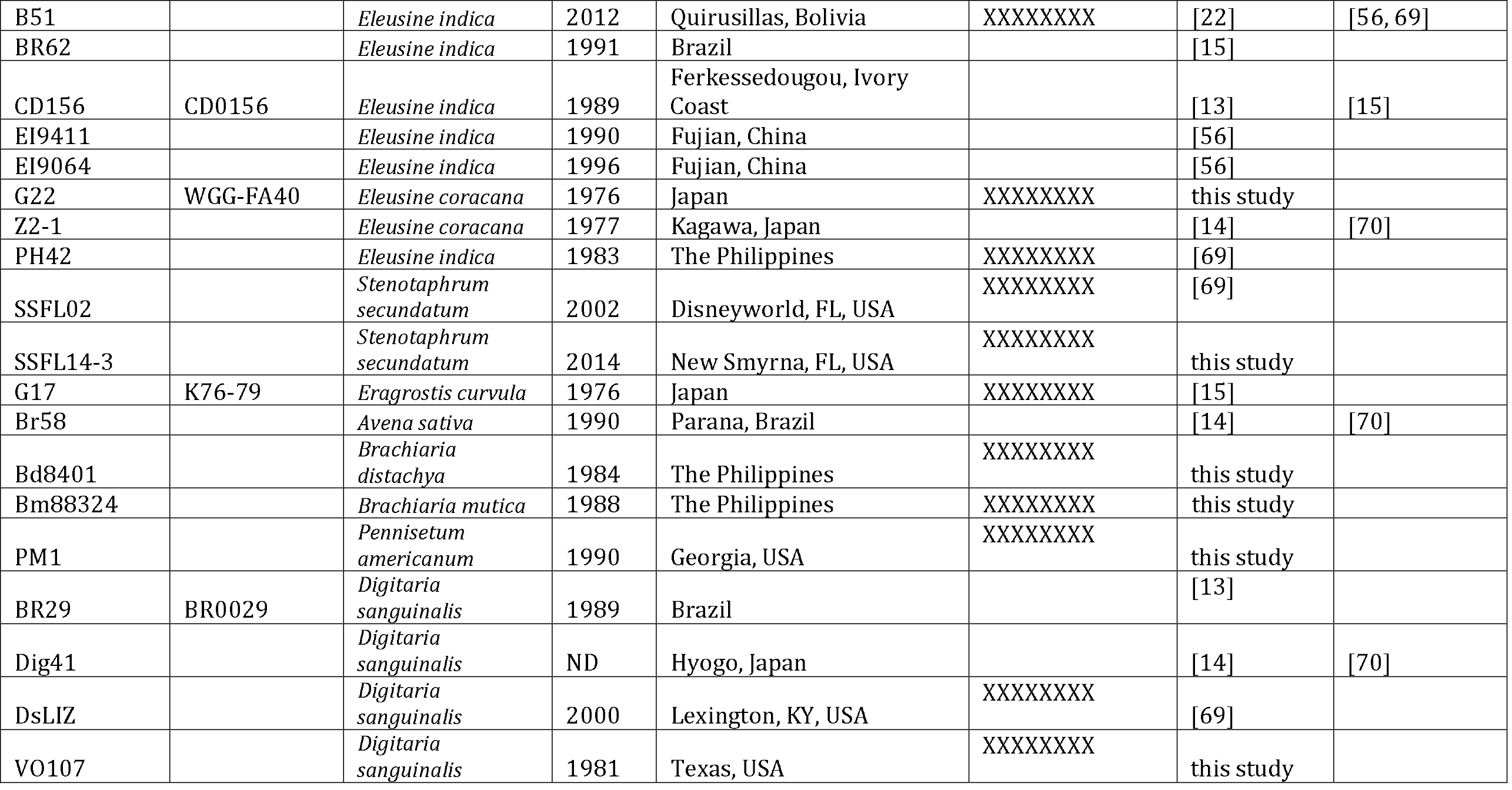
Magnaporthe oryzae, M. grisea and M. pennisetigena strains used in this study.

| Isolate ID | Synonym | Host | Year | Locality | NCBI accession | Sequence source | References |
| --- | --- | --- | --- | --- | --- | --- | --- |
| BdBar | BdBar16-1 | *Triticum aestivum* | 2016 | Barisal, Bangladesh |  | [23] |  |
| BdJes | BdJes16-1 | *Triticum aestivum* | 2016 | Jessore, Bangladesh |  | [23] |  |
| BdMeh | BdMeh16-1 | *Triticum aestivum* | 2016 | Mehepur, Bangladesh |  | [23] |  |
| B2 |  | *Triticum aestivum* | 2011 | Bolivia | XXXXXXXX | [69] |  |
| B71 |  | *Triticum aestivum* | 2012 | Bolivia |  | [23] | [69] |
| Br7 |  | *Triticum aestivum* | 1990 | Parana, Brazil | XXXXXXXX | [22] | [69] |
| BR0032 | BR32 | *Triticum aestivum* | 1991 | Brazil |  | [13] | [15] |
| Br48 |  | *Triticum aestivum* | 1990 | Mato Grosso do Sul, Brazil |  | [14] | [22] |
| Br80 |  | *Triticum aestivum* | 1991 | Brazil | XXXXXXXX | [22] | [69] |
| Br130 |  | *Triticum aestivum* | 1990 | Mato Grosso do Sul, Brazil | XXXXXXXX | [22] |  |
| P3 |  | *Triticum durum* | 2012 | Canindeyu, Paraguay | XXXXXXXX | [69] |  |
| PY0925 |  | *Triticum aestivum* | 2009 | Predizes, Brazil |  | [15] |  |
| PY36-1 | PY36.1 | *Triticum aestivum* | 2007 | Brasilia, Brazil |  | [15] |  |
| PY5003 | PY05003 | *Triticum aestivum* | 2005 | Londrina, Brazil |  | [15] |  |
| PY5010 | PY05010 | *Triticum aestivum* | 2005 | Londrina, Brazil |  | [15] | [69] |
| PY5033 | PY05033 | *Triticum aestivum* | 2005 | Londrina, Brazil |  | [15] |  |
| PY6017 | PY06017 | *Triticum aestivum* | 2006 | Coromandel, Brazil |  | [15] |  |
| PY6045 | PY06045 | *Triticum aestivum* | 2006 | Goiania, Brazil |  | [15] |  |
| PY86-1 | PY86.1 | *Triticum aestivum* | 2008 | Cascavel, Brazil |  | [15] |  |
| T25 |  | *Triticum aestivum* | 1988 | Parana, Brazil | XXXXXXXX | [22] | [69] |
| WHTQ |  | *Triticum aestivum* | ND | Brazil | XXXXXXXX | [22] |  |
| WBKY11 | WBKY11-15 | *Triticum aestivum* | 2011 | Lexington, KY, USA | XXXXXXXX | [22] | [69] |
| P28 | P-0028 | *Bromus tectorum* | 2014 | Paraguay | XXXXXXXX | [69] |  |
| P29 | P-0029 | *Bromus tectorum* | 2014 | Paraguay | XXXXXXXX | [69] |  |
| CHRF |  | *Lolium perenne* | 1996 | Silver Spring, MD, USA | XXXXXXXX | [69] |  |
| CHW |  | *Lolium perenne* | 1996 | Annapolis, MD, USA | XXXXXXXX | [69] |  |
| FH |  | *Lolium perenne* | 1997 | Hagerstown, MD, USA | XXXXXXXX | [22] | [69] |
| GG11 |  | *Lolium perenne* | 1997 | Lexington, KY, USA | XXXXXXXX | [22] |  |
| HO |  | *Lolium perenne* | 1996 | Richmond, PA, USA | XXXXXXXX | [22] |  |
| LpKY97 | LpKY97-1 | *Lolium perenne* | 1997 | Lexington, KY, USA | XXXXXXXX | [22] | [69] |
| PgKY | PgKY4OV2.1 | *Lolium perenne* | 2000 | Lexington, KY, USA |  | [15] |  |
| PGPA | PgPA18C-02,<br>PgPA | *Lolium perenne* | 1998 | Pennsylvania, USA |  | [15] |  |
| PL2-1 |  | *Lolium multiflorum* | 2002 | Pulaski Co., KY, USA | XXXXXXXXX | [22] |  |
| PL3-1 |  | *Lolium multiflorum* | 2002 | Pulaski Co., KY, USA | XXXXXXXXX | [22] | [69] |
| Pg1213-22 |  | *Festuca*<br>*arundinaceum* | 1999/<br>2000 | GA | XXXXXXXXX | [69] |  |
| TF05-1 |  | *Festuca*<br>*arundinaceum* | 2005 | Lexington, KY, USA | XXXXXXXXX | this study |  |
| IB33 |  | *Oryza sativa* | ND | Texas, USA | XXXXXXXXX | A. Marchetti |  |
| FR13 | FR0013 | *Oryza sativa* | 1988 | France |  | [13] | [15] |
| GY11 | GY0011,<br>Guy11 | *Oryza sativa* | 1988 | French Guyana |  | [13] | [15, 22] |
| IA1 | ARB114 | *Oryza sativa* | 2009 | Arkansas, USA | XXXXXXXXX | [69] |  |
| IB49 | ZN61 | *Oryza sativa* | 1992 | Arkansas, USA | XXXXXXXXX | [69] |  |
| IC17 | ZN57 | *Oryza sativa* | 1992 | Arkansas, USA | XXXXXXXXX | [69] |  |
| IE1K | TM2 | *Oryza sativa* | 2003 | Arkansas, USA | XXXXXXXXX | [69] |  |
| INA168 | Ina168 | *Oryza sativa* | 1958 | Aichi, Japan |  | [14] | [70] |
| KEN53-33 | Ken53-33 | *Oryza sativa* | 1953 | Aichi, Japan |  | [14] |  |
| ML33 |  | *Oryza sativa* | 1995 | Mali | XXXXXXXXX | this study |  |
| P131 |  | *Oryza sativa* | ND | Japan |  | [55] | [22, 56] |
| Y34 |  | *Oryza sativa* | 1982 | China, Yunnan |  | [55] | [22, 56] |
| P-2 | P2 | *Oryza sativa* | 1948 | Aichi, Japan |  | [14] |  |
| PH0014-rn | PH0014,<br>PH14 | *Oryza sativa* | ND | The Philippines |  | [13] | [15] |
| TH3 |  | *Oryza sativa* | ND | Thailand |  | [14] | [14] |
| 87-120 |  | *Oryza sativa* | ND |  | XXXXXXXXX | this study |  |
| TH0012-rn | TH0012,<br>TH12 | *Hordeum vulgare* | ND | Thailand |  | [13] | [15] |
| TH0016 | TH16 | *Hordeum vulgare* | ND | Thailand |  | [13] | [15] |
| Arcadia |  | *Setaria viridis* | 1998 | Lexington, KY, USA | XXXXXXXXX | [22] | [69] |
| US0071 | US71 | *Setaria spp.* | ND | USA |  | [13] | [15] |
| GrF52 |  | *Setaria viridis* | 2001 | Lexington, KY, USA | XXXXXXXXX | this study |  |
| KANSV1-4-1 | KNSV | *Setaria viridis* | 1975 | Kanagawa, Japan |  | [14] |  |
| SA05-43 |  | *Setaria viridis* | 2005 | Nagasaki, Japan |  | [14] |  |
| Sv9610 |  | *Setaria viridis* | 1996 | Zhejiang, China |  | [56] |  |
| Sv9623 |  | *Setaria viridis* | 1996 | Zhejiang, China |  | [56] |  |
| GFSI1-7-2 | GFSI | *Setaria italica* | 1977 | Gifu, Japan |  | [14] |  |
| B51 |  | *Eleusine indica* | 2012 | Quirusillas, Bolivia | XXXXXXXX | [22] | [56, 69] |
| BR62 |  | *Eleusine indica* | 1991 | Brazil |  | [15] |  |
| CD156 | CD0156 | *Eleusine indica* | 1989 | Ferkessedougou, Ivory Coast |  | [13] | [15] |
| EI9411 |  | *Eleusine indica* | 1990 | Fujian, China |  | [56] |  |
| EI9064 |  | *Eleusine indica* | 1996 | Fujian, China |  | [56] |  |
| G22 | WGG-FA40 | *Eleusine coracana* | 1976 | Japan | XXXXXXXX | this study |  |
| Z2-1 |  | *Eleusine coracana* | 1977 | Kagawa, Japan |  | [14] | [70] |
| PH42 |  | *Eleusine indica* | 1983 | The Philippines | XXXXXXXX | [69] |  |
| SSFL02 |  | *Stenotaphrum secundatum* | 2002 | Disneyworld, FL, USA | XXXXXXXX | [69] |  |
| SSFL14-3 |  | *Stenotaphrum secundatum* | 2014 | New Smyrna, FL, USA | XXXXXXXX | this study |  |
| G17 | K76-79 | *Eragrostis curvula* | 1976 | Japan | XXXXXXXX | [15] |  |
| Br58 |  | *Avena sativa* | 1990 | Parana, Brazil |  | [14] | [70] |
| Bd8401 |  | *Brachiaria distachya* | 1984 | The Philippines | XXXXXXXX | this study |  |
| Bm88324 |  | *Brachiaria mutica* | 1988 | The Philippines | XXXXXXXX | this study |  |
| PM1 |  | *Pennisetum americanum* | 1990 | Georgia, USA | XXXXXXXX | this study |  |
| BR29 | BR0029 | *Digitaria sanguinalis* | 1989 | Brazil |  | [13] |  |
| Dig41 |  | *Digitaria sanguinalis* | ND | Hyogo, Japan |  | [14] | [70] |
| DsLIZ |  | *Digitaria sanguinalis* | 2000 | Lexington, KY, USA | XXXXXXXX | [69] |  |
| VO107 |  | *Digitaria sanguinalis* | 1981 | Texas, USA | XXXXXXXX | this study |  |

ND, no data. “References” lists studies that used the sequencing data, besides present study. Isolates Br116.5, Br118.2, TP2, MZ5-1-6, and Br35 sequenced by Inoue et al. [70], Bangladeshi isolates and isolates PY05002, PY06025, PY06047, PY25.1, PY35.3, PY05035 sequenced by Islam et al. [15], isolate SA05-144 sequenced by Yoshida et al. [14], and isolates DS9461 and DS0505 sequenced by Zhong et al. [56] were not included in the study.

First we employed the multivariate approach implemented in Discriminant Analysis of Principal Components (DAPC; [30]) to examine population subdivision within *M. oryzae*. Using the haplotypes identified from orthologous loci, the Bayesian Information Criterion plateaued at K=10 in models varying K from 2 to 20 clusters, indicating that K=10 captures the most salient features of population subdivision (Figure S3). Clusters identified at K=10 were as follows: (1) isolates from rice and two isolates from barley (dark green; referred to as the *Oryza* lineage); (2) isolates from *Setaria* sp. (light green; referred to as the *Setaria* lineage); (3) isolate Bm88324 from *Brachiaria mutica* (olive; referred to as the *Brachiaria1 lineage*); (4) isolate Bd8401 from *Brachiaria distachya* (brown; referred to as the *Brachiaria2* lineage); (5) isolates from *Stenotaphrum* (red; referred to as the *Stenotaphrum* lineage); (6) 17 of the 22 isolates from wheat and an isolate from *Bromus* (blue; referred to as the *Triticum lineage*); (7) the remaining 3/22 isolates from wheat together with isolates from *Lolium, Festuca*, oat and a second isolate from *Bromus* (purple; referred to as the *Lolium* lineage); (8 & 9) isolates from *Eleusine* that formed two distinct clusters (light orange and orange; referred to as the *Eleusine1* and *Eleusine2* lineages, respectively); and (10) an isolate from *Eragrostis* (yellow; referred to as the *Eragrostis lineage*) (Figure 1). Increasing K mostly resulted in further subdivision among the isolates from wheat, rice and *Lolium* sp. The discovery of three wheat blast isolates that grouped with the *Festuca-Lolium* pathogens was important because it supports the idea that wheat-infecting isolates belong to at least two distinct populations.

**Figure 1.**
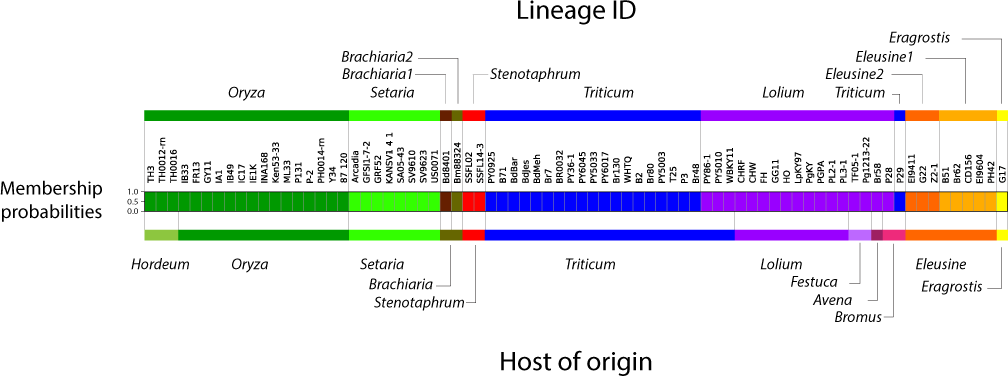
Discriminant Analysis of Principal Components, assuming K=10 clusters. Each isolate is represented by a thick vertical line divided into K segments that represent the isolate’s estimated membership probabilities in the K=10 clusters (note that all isolates have high membership probabilities in a single cluster, hence only a single segment is visible). The host of origin of samples is shown below the barplot, and lineage IDs are shown above the barplot.

Next, we inferred gene genealogies using maximum-likelihood and distance-based methods. Both approaches produced trees that corresponded well with the subdivisions identified in DAPC. The tree generated using maximum likelihood (ML) analysis of orthologous genes displayed a topology with ten lineages (Figure 2) showing one-to-one correspondence with the K clusters from DAPC (Figure 1; Figure S3). Nine of these lineages had >90% bootstrap support. The lineage that corresponded to the “blue” DAPC cluster (including the 17 isolates from wheat and isolate P29 from *Bromus*) had poor bootstrap support (50%).

**Figure 2.**
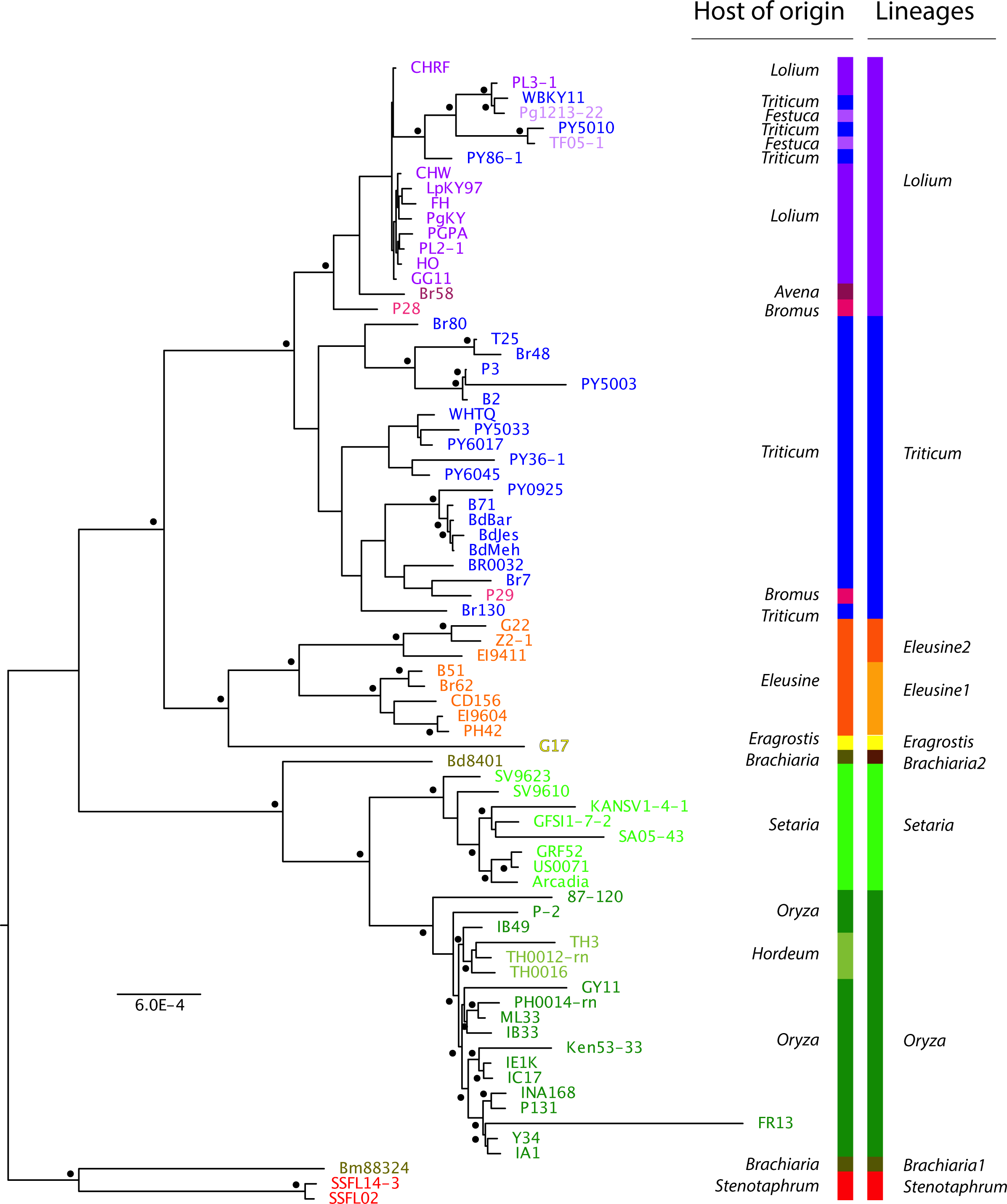
Maximum likelihood tree based on the concatenation of 2,682 orthologous coding sequences extracted from 76 *M. oryzae genome*. Nodes with bootstrap support >90% are indicated by dots (100 bootstrap replicates).

The neighbor-joining tree built using “total genome” pairwise distances resolved very similar groupings to the DAPC (Figure 1; Figure S3) and ML ortholog tree (Figure 3). The only major discrepancy between ML and NJ trees was the confident placement of 87-120 – an isolate from rice – outside of the rice clade in the NJ tree (Figure 3).

**Figure 3.**
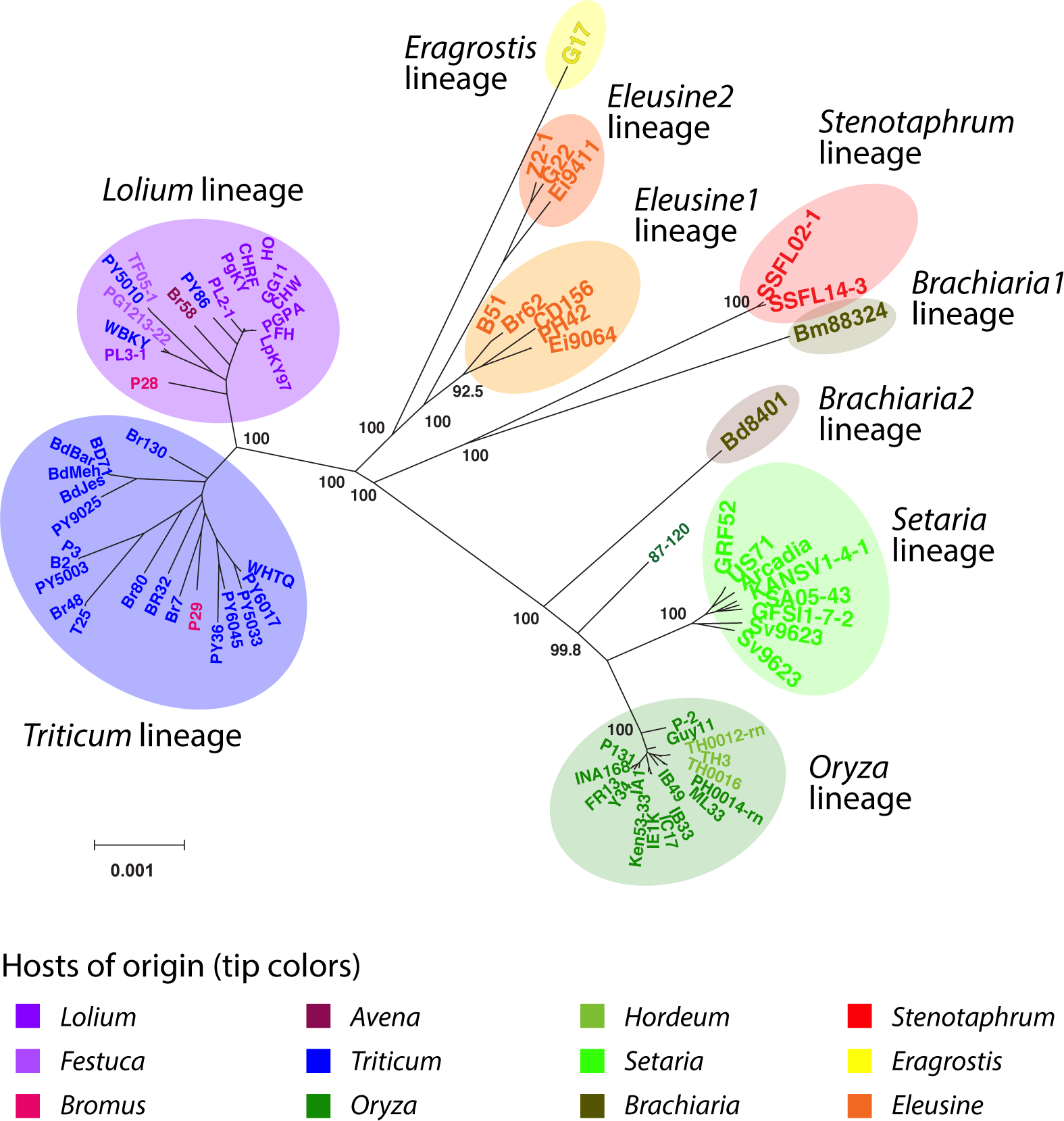
Total evidence neighbor-joining distance tree using pairwise distances (number of differences/kb) calculated from analysis of pairwise blast alignments between repeat-masked genomes. Only nodes with confidence > 80% (see methods) are labeled. Gray ovals are drawn around the main host-specialized populations for clarity.

### Levels of polymorphism within and divergence between lineages/species

We compared levels of polymorphism within lineages to levels of divergence between lineages or species to apprehend the relative evolutionary depth of the lineages within *M. oryzae*. Genetic variability based on 2,682 orthologs was relatively low and one order of magnitude higher in the rice and wheat lineages (0.1% differences per site) than in the Lolium and *Setaria* lineages (other lineages not included in the calculations due to small sample sizes – only lineages with n>6 included; Table 2). The null hypothesis of no recombination could be rejected in the *Lolium*, wheat, rice and *Setaria* lineages using the Pairwise Homoplasy Test implemented in the SPLITSTREE 4.13 program ([31]; p-value:0.0; Table 2).

**Table 2.**
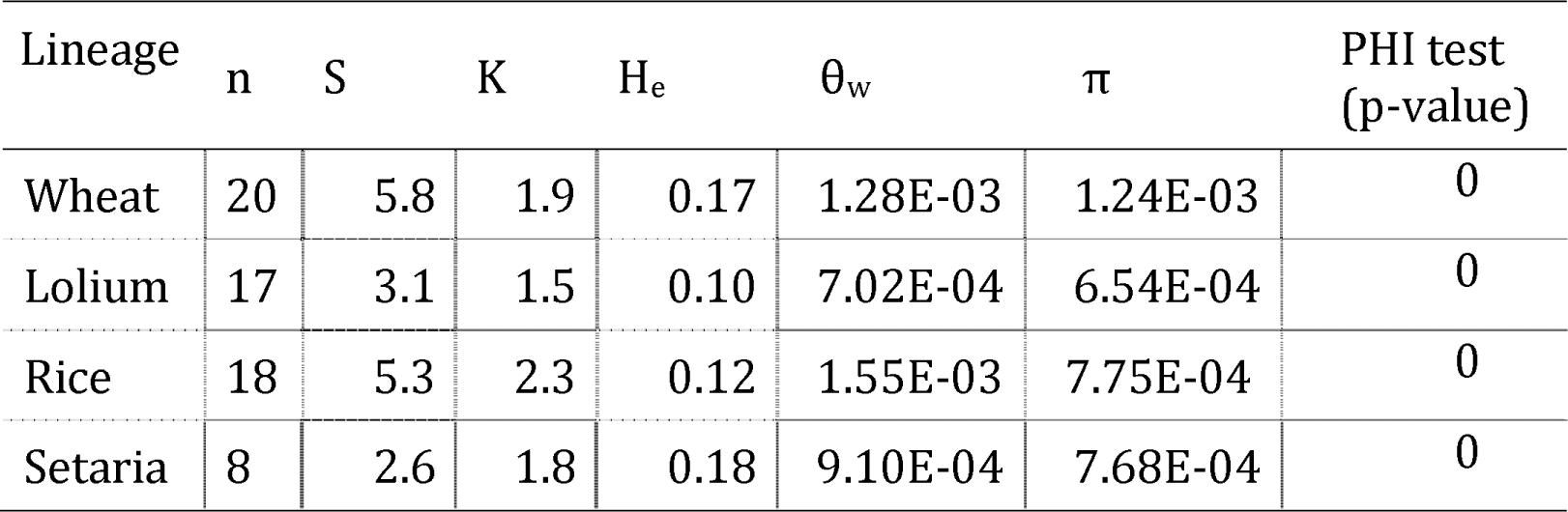
Summary of population genetic variation at 2682 single-copy orthologous genes in wheat, lolium, rice and setaria lineages of *Magnaporthe oryzae*.

| Lineage | n | S | K | H<sub>e</sub> | θ<sub>w</sub> | π | PHI test (p-value) |
| --- | --- | --- | --- | --- | --- | --- | --- |
| Wheat | 20 | 5.8 | 1.9 | 0.17 | 1.28E-03 | 1.24E-03 | 0 |
| Lolium | 17 | 3.1 | 1.5 | 0.10 | 7.02E-04 | 6.54E-04 | 0 |
| Rice | 18 | 5.3 | 2.3 | 0.12 | 1.55E-03 | 7.75E-04 | 0 |
| Setaria | 8 | 2.6 | 1.8 | 0.18 | 9.10E-04 | 7.68E-04 | 0 |

Other lineages were not included in calculations because of too small a sample size (n<6); n is sample size; θ_w_ is Watterson’s θ per bp; π is nucleotide diversity per bp; H_e_ is haplotype diversity; K is the number of haplotypes. PHI test is the Pairwise Homoplasy Test. The PHI test is implemented in SPLITSTREE. The null hypothesis of no recombination was tested, for the PHI test using random permutations of the positions of the SNPs based on the expectation that sites are exchangeable if there is no recombination

Genome-wide nucleotide divergence was one order of magnitude higher between *M. oryzae* and its closest relatives, *M. grisea* and *M. pennisetigena*, than it was among isolates within *M. oryzae*. The maximum pairwise distance (number of differences per kilobase) between any two *M. oryzae* isolates was less than 1%, genome-wide (Figure S4; Table S1), compared with *M. oryzae* vs *M. grisea*, *M. oryzae* vs. *M. pennisetigena*, or *M. grisea* vs *M. pennisetigena*, all of which were consistently greater than 10%. The low level of genetic divergence among *M. oryzae* isolates, compared with that observed when comparing *M. oryzae* isolates to other established related species, provides good evidence against the existence of relatively ancient cryptic species within *M. oryzae.* (Table S1).

### Re-assessment of Pgt as a novel species using whole genome data

While the 10 loci utilized in the Castroagudin et al. [24] study do not support the *Pgt* split based on GCPSR criteria, our DAPC and whole genome ML and NJ analyses supported the partitioning of wheat blast isolates into two, genetically-distinct lineages: one consisting almost exclusively of wheat-infecting isolates, the other comprising largely *Festuca*‐ and *Lolium*-infecting isolates as well as few wheat-infecting isolates (Figure 2; Figure 3). However, the Castroagudin et al. study did not include *Festuca*‐ and *Lolium*-infecting isolates and genome sequences from this study are not available. Therefore, to test for possible correspondence between the proposed *Pgt* species and the *Lolium* lineage (or indeed the *Triticum* lineage), we extended the 10 loci analysis to the *M. oryzae* genome sequences used in the present study. For reference, we included the multilocus data for 16 isolates from the Castroagudin et al., representing all the major clades from that study. Nine of the 10 loci were successfully recovered from 68 of our *M. oryzae* genome sequences. The remaining locus, CH7-BAC9, was absent from too many genome sequences and, as a result, was excluded from the analysis.

The nine concatenated loci produced a total-evidence RAxML tree in which very few branches had bootstrap support greater than 50% (Figure 4). All of the *Pgt* isolates from the Castroagudin et al. study were contained in a clade with 80% support. Inspection of the *MPG1* marker that was reported to be diagnostic for *Pgt* (Castroagudin et al. 2016) revealed that all of the isolates in this clade contained the *Pgt*-type allele (green dots) and should therefore be classified as *Pgt* (Figure 4). Critically, however, a few isolates outside this clade also harbored the *Pgt*-type allele. Moreover, the clade also included isolates from the present study which came from wheat, annual ryegrass, perennial ryegrass, tall fescue, finger millet, and goosegrass – isolates that did not group together in the DAPC analysis (Figure 1), or in the ML and NJ trees built using the orthologous genes or whole genome SNP data (Figure 2; Figure 3). Isolates carrying *Pgt*-type allele were in fact distributed among three genetically distinct and well-supported clades (Figure 2; Figure 3). Furthermore, visual inspection of the topologies and bootstrap supports for each single-locus tree revealed that GCPSR criteria were not satisfied for the clade including all of the *Pgt* isolates from the Castroagudin et al. Thus, isolates characterized by Castroagudin et al. [24] as *Pgt* fail to constitute a phylogenetically cohesive group based on total genome evidence and, thus, the existence of the *Pgt* species is not supported by our new genome-wide data and analyses.

**Figure 4.**
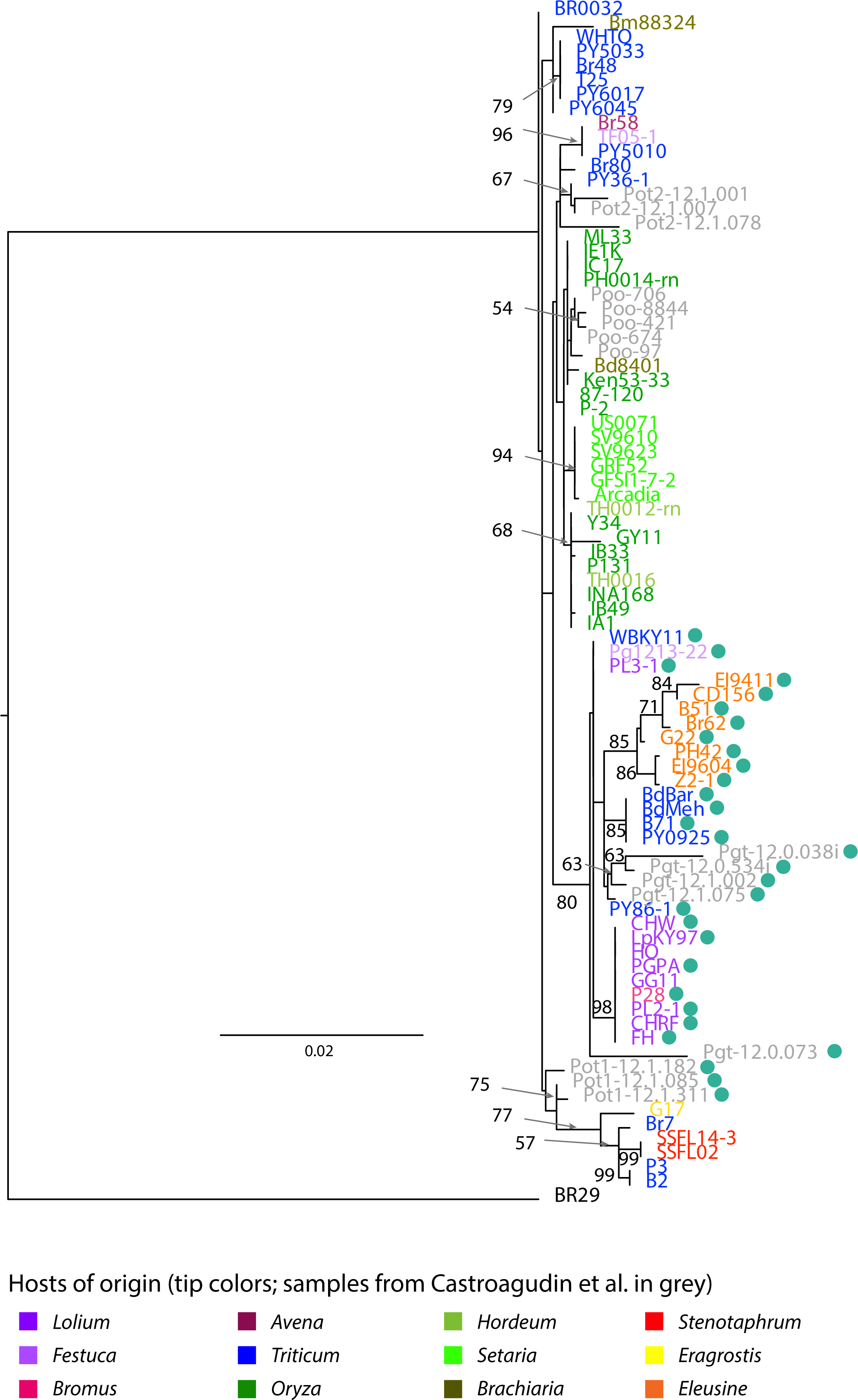
Maximum likelihood tree based on concatenated dataset comprising nine loci used in Castroagudin et al. [24], retrieved from 76 *M. oryzae* genomes. Numbers above branches represent bootstrap supports after 100 bootstrap replicates. Only nodes with bootstrap support > 50 are labelled. Representatives of isolates used by Castroagudin et al. [24] in their study were included in the analysis and are colored in light grey. Green dots mark the strains containing the *Pgt*-type allele according to Castroagudin et al. [24].

The basis for the previous designation of *Pgt* as a novel species was clearly revealed when *MPG1* alleles were mapped onto the ML and NJ trees. The distribution of *MPG1* alleles among different *M. oryzae* lineages was discontinuous (Figure S5). As an example, isolates from the *Triticum* lineage carried three different *MPG1* alleles. Two of these (including the *Pgt*-type) were also present in the *Lolium* lineage, while the third *MPG1* (ACT17T-C-6CAA140, Figure S5) was shared by distantly-related isolates from the Stenotaphrum lineage (Figure S5). Isolates from the *Eleusine* lineage also carried *Pgt*-type *MPG1* allele and two other variants, while isolates from the *Setaria* and *Oryza* lineages carried an *MPG1* allele distinct from all the others (Figure S5). Overall, the distribution of *MPG1* alleles points to the occurrence of incomplete lineage sorting and gene flow during *M. oryzae* diversification. Importantly, seven markers studied by Castroagudin et al. – including *MPG1* - showed discontinuities in their distributions among lineages defined using genome-wide data and analyses (Figure S5). The two other markers (ACT1 and CHS1) used by Castroagudin et al. showed no sequence variations among the 68 *M. oryzae* isolates analyzed in the present study (data not shown) and are not useful for phylogenetic classification.

### Species tree inference and phylogenetic species recognition from genome-wide data

The total evidence genealogies generated using sequence data from 76 *M. oryzae* genomes using either distance-based (whole genomes) or maximum likelihood (2,682 single-copy orthologs) phylogenetic methods were highly concordant in terms of lineage composition and branching order (Figure 2; Figure 3). However, concatenation methods can be positively misleading, as they assume that all gene trees are identical in topology and branch lengths and they do not explicitly model the relationship between the species tree and gene trees [32]. To estimate the “species tree” and to re-assess previous findings of cryptic species within *M. oryzae*, we used a combination of species inference using the multispecies coalescent method implemented in ASTRAL [27-29] and a new implementation of the GCPSR that can handle genomic data.

The ASTRAL “species tree” with the local q1 support values on key branches is shown in Figure 5. The four *M. grisea* isolates from crabgrass (*Digitaria* sp.) and the *M. pennisetigena* isolate from fountaingrass (*Pennisetum* sp.) were included as outgroups, bringing the total number of isolates to 81 and reducing the dataset to 2,241 single-copy orthologous genes. The branches holding the clades containing the wheat blast isolates had q1 support values of 0.49, 0.39 and 0.37 which means that, in each case, fewer than 50% of the whole set of quartet gene trees recovered from the individual gene genealogies agreed with the local topology around these branches in the species tree. The branches that separated *M. grisea* and *M. pennisetigena* from *M. oryzae* had respective q1 values of 1, providing strong support for relatively ancient speciation. In contrast, the highest q1 value on any of the branches leading to the host-specialized clades was 0.8 for the *Setaria* pathogens, indicating that approximately 20% of the quartets recovered from individual gene trees were in conflict with the species tree around this branch. Together, these results indicate high levels of incomplete lineage sorting within, and/or gene flow involving these groups, and are thus inconsistent with the presence of genetically isolated lineages (i.e. species).

**Figure 5.**
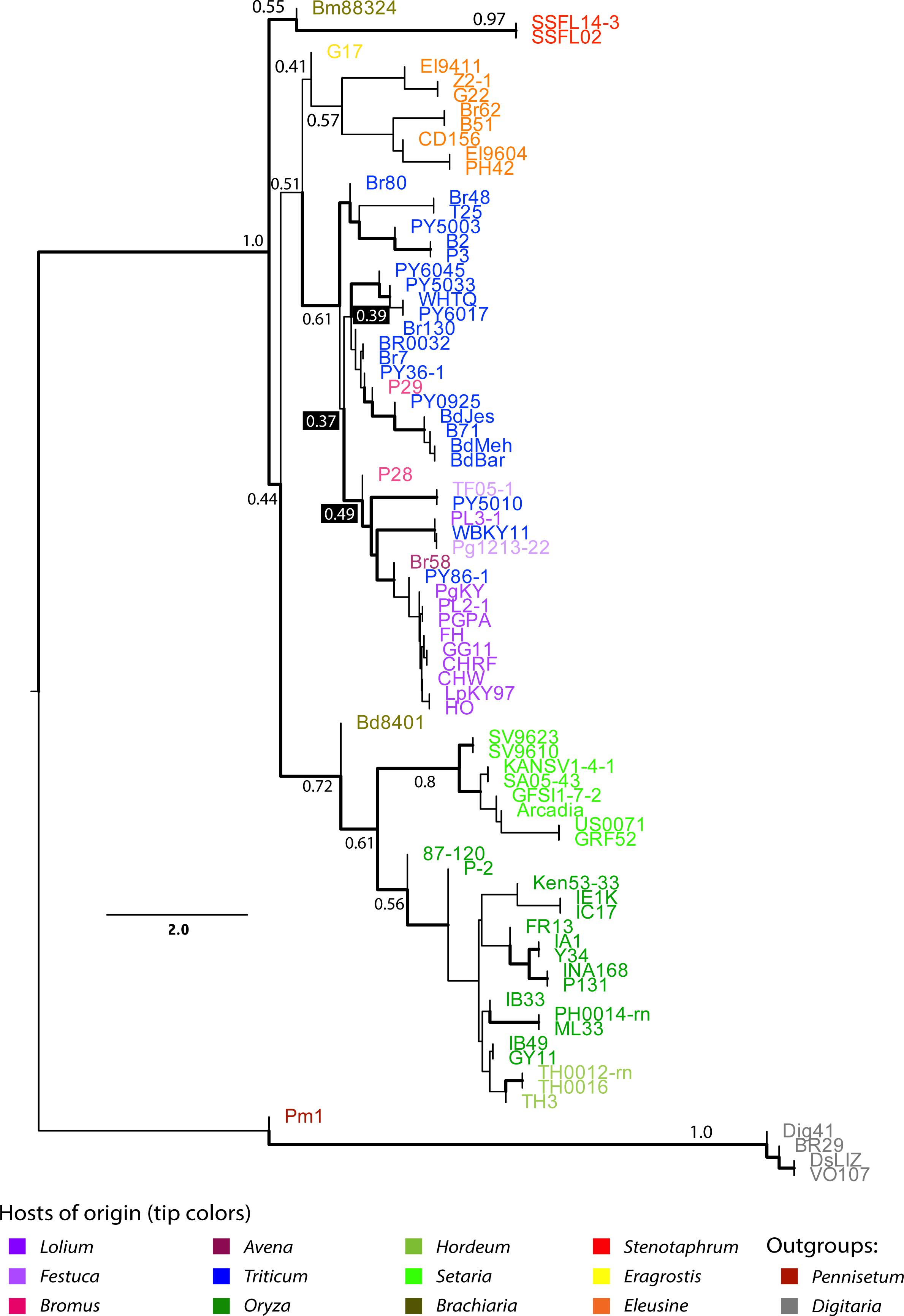
ASTRAL analysis to test for incomplete lineage sorting/gene flow among 81 Magnaporthe genomes, using 2,241 single-copy orthologous sequence loci. Thicker branches represent branches that have a bootstrap support > 50 after multilocus bootstrapping. Number above branches represent q1 local support (i.e. the proportion of quartet trees in individual genealogies that agree with the topology recovered by the ASTRAL analysis around the branch), with q1 values showed on black background for branches holding wheat blast isolates.

As a formal test for the presence of cryptic species within *M. oryzae*, we applied the phylogenetic species recognition criteria to the set of 2,241 single-copy orthologous genes using an implementation of the GCPSR scalable to any number of loci. Applying the GCPSR following the non-discordance criterion of Dettman et al. (a clade has to be well supported by at least one single-locus genealogy and not contradicted by any other genealogy at the same level of support; [25]) resulted in the recognition of no species within *M. oryzae.*

### Historical gene flow between lineages

The existence of gene flow and/or incomplete lineage sorting was also supported by phylogenetic network analysis. We used the network approach neighbor-net implemented in SPLITSTREE 4.13 [25] to visualize evolutionary relationships, while taking into account the possibility of recombination within or between lineages. The network inferred from haplotypes identified using the 2,682 single-copy orthologs in the 76 *M. oryzae* strains, showed extensive reticulation connecting all lineages, consistent with recombination or incomplete lineage sorting (Figure 6).

**Figure 6.**
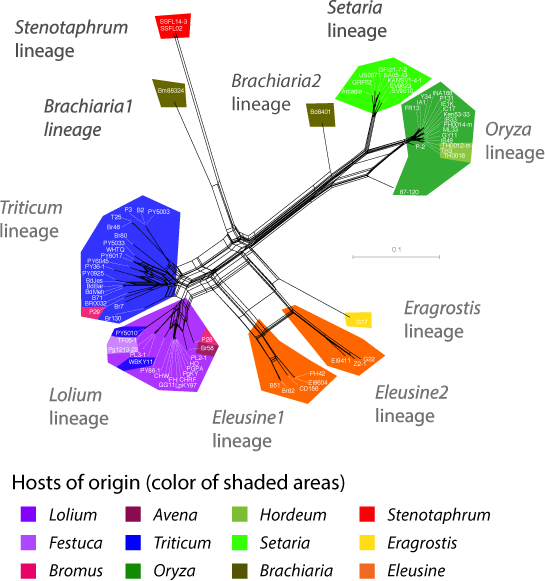
Neighbor-Net network built with SPLITSTREE. The figure shows relationships between haplotypes identified based on the full set of 25,078 SNPs identified in 2,682 single-copy orthologs, excluding sites missing data, gaps and singletons.

To disentangle the role of gene flow versus incomplete lineage sorting in observed network reticulations, but also to gain insight into the timing and extent of genetic exchanges, we used ABBA/BABA tests, which compare numbers of two classes of shared derived alleles (the ABBA and BABA classes). For three lineages P1, P2 and P3 and an outgroup with genealogical relationships (((P1,P2),P3),O), and under conditions of no gene flow, shared derived alleles between P2 and P3 (ABBA alleles) and shared derived alleles between P1 and P3 (BABA alleles) can only be produced by incomplete lineage sorting, and should be equally infrequent [33]. Differences in numbers of ABBA and BABA alleles are interpreted as gene flow assuming no recurrent mutation and no deep ancestral population structure within lineages. We computed D, which measures the imbalance between numbers of ABBA and BABA sites and is used to test for admixture in ((P1,P2),P3) triplets, with D>0 implying gene flow between P2 and P3, and D<0 implying gene flow between P1 and P3 [34, 35]. We also made use of the heterogeneity in divergence time between members of ((P1,P2),P3) triplets to examine gene flow across three time periods [33], following these principles: (i) triplets including the most recently diverged lineages as P1 and P2 (i.e. the *Triticum* and *Lolium* lineages, the two *Eleusine* lineages, or the *Oryza* and *Setaria* lineages) carried information about gene flow across relatively recent times, (ii) triplets including, as P1 and P2, two lineages from the same main group of lineages (i.e. *Eragrostis/Eleusine1/Eleusine2/Triticum/Lolium* or *Brachiaria2/Setaria/Oryza*, excluding (P1,P2) pairs already used in (i)) carried information about gene flow across intermediate times, and (iii) triplets including, as P1 and P2, two lineages from different main groups of lineages (i.e. *Eragrostis / Eleusine1 / Eleusine2 / Triticum / Lolium* and *Brachiaria2 / Setaria / Oryza*) and *Stenotaphrum* or *Brachiaria1* as P3 carried information about gene flow across a relatively long time period (Figure S6).

The D statistic measuring differences in counts of ABBA and BABA alleles was significantly different from zero (Z-score>3) in 104 of 120 lineage triplets, consistent with a history of gene flow between lineages within *M. oryzae* (Table S2). Given that a (P1,P2) pair can be represented as multiple ((P1,P2),P3) triplets, and that the sign of D indicates what is the pair involved in gene flow within each triplet, the 104 triplets with significant D values in fact represented 35 pairs connected by gene flow, spanning the three time scales defined by the phylogenetic affiliation of lineages (Figure S6). Lineages were equally represented in triplets deviating from null expectations assuming no gene flow, no ancient structure and no recurrent mutations. Consistent with historical gene flow, searches for private allele found no gene, among the 2241 gene surveyed, carrying mutations exclusive to a single lineage. Together, these results indicate that gene flow was widespread, both across historical times and lineages, but it cannot be excluded that much of the signal was caused by events that happened prior to lineage splitting.

### Recent admixture and gene flow between lineages

We then used the program STRUCTURE [36-38] to detect possible recent admixture between lineages (Figure S3). STRUCTURE uses Markov chain Monte Carlo simulations to infer the assignment of genotypes into K distinct clusters, minimizing deviations from Hardy– Weinberg and linkage disequilibria within each cluster. The patterns of clustering inferred with STRUCTURE were largely similar to those inferred with DAPC. STRUCTURE analysis provided evidence for admixture at all K values (Figure S3), suggesting that recent admixture events have recently shaped patterns of population subdivision within *M. oryzae.* ‘Chromosome painting’, a probabilistic method for reconstructing the chromosomes of each individual sample as a combination of all other homologous sequences [39], also supported the lack of strict genetic isolation between lineages (Text S1).

## Discussion

### Population subdivision but no cryptic phylogenetic species

Using population- and phylogenomic analyses of single-copy orthologous genes and whole-genome SNPs identified in *M. oryzae* genomes from multiple cereal and grass hosts, we provide evidence that *M. oryzae* is subdivided in multiple lineages preferentially associated with one host plant genus. Neither the re-analysis of previous data, nor the analysis of new data using previous phylogenetic species recognition markers, supports the existence of a wheat blast-associated species called *P. graminis-tritici* [24]. Marker *MPG1*, which holds most of the divergence previously detected, does not stand as a diagnostic marker of the wheat-infecting lineage of *M. oryzae* when tested in other lineages. Previous conclusions about the existence of cryptic species *P. graminis-tritici* also stem from the fact that available information on *M. oryzae* diversity had been insufficiently taken into account. In particular, isolates from the lineages most closely related to wheat strains (i.e. isolates from the *Lolium* lineage; [11, 12, 15, 22]) were not represented in previous species identification work [24]. Using phylogenetic species recognition by genealogical concordance we could not identify cryptic phylogenetic species and thus *M. oryzae* is not, strictly speaking, a species complex. As a consequence, *Pyricularia graminis-tritici* cannot - and should not – be considered as a valid name for wheat-infecting strains, because (1) it refers to a subset of wheat-infecting strains, and quarantine on *P. graminis-tritici* alone would not prevent introduction of aggressive wheat blast pathogens (2) it groups very aggressive wheat pathogens from South America and South Asia with *Eleusine*-infecting strains that are largely distributed in the world. Given the devastating potential of wheat blast disease, it is vital that accurate strain identification and species assignment can be carried out by plant health agencies in order to safeguard against importation and spread of the disease. Correct species assignment is therefore a critical consideration. Hence, although the formal rules of taxonomy would imply treating *P. graminis-tritici* as synonym of *Magnaporthe oryzae*, we strongly recommend dismissal of *P. graminis-tritici* as a valid name to refer to wheat-infecting strains of *M. oryzae.*

### Incipient speciation by ecological specialization following host-shifts

Several features of the life cycle of *M. oryzae* are conducive to speciation by ecological specialization following host shifts, suggesting that the observed pattern of population subdivision in *M. oryzae* actually corresponds to ongoing speciation. Previous experimental measurements of perithecia formation and ascospore production – two important components of reproductive success-suggested inter-fertility in vitro between most pairs of lineages with high levels of ascospore viability [40-43]. This suggests that intrinsic pre- or post-mating reproductive barriers, such as assortative mating by mate choice or gametic incompatibility, and zygotic mortality, are not responsible for the relative reproductive isolation between lineages – which creates the observed pattern of population subdivision. Instead, the relative reproductive isolation between lineages could be caused by one or several pre‐ or post-mating barriers (Table 1 in [44]), such as mating-system isolation or hybrid sterility (intrinsic barrier), or difference in mating times, difference in mating sites, immigrant inviability, or ecologically-based hybrid inviability (extrinsic barriers).

Previous pathogenicity assays revealed extensive variability in the host range of *M. oryzae* isolates, and both in terms of pathogenicity towards a set of host species or pathogenicity towards a set of genotypes from a given host [41, 45]. Indeed, extensive genetic analyses show that host species specificity in *M. oryzae*, similar to rice cultivar specificity, could be controlled by a gene-for-gene relationship in which one to three avirulence genes in the fungus prevent infection of particular host species [43, 46, 47]. Loss of the avirulence genes would allow infection of novel hosts to occur. Additionally, host species specificity is not strictly maintained. Under controlled conditions, most lineages have at least one host in common [45], and strains within one lineage can still cause rare susceptible lesions on naive hosts [21, 48]. Moreover, a single plant infected by a single genotype can produce large numbers of spores in a single growing season [49], allowing the pathogen to persist on alternative host even if selection is strong, and promoting the rapid and repeated creation of genetic variation [6]. Although some of these features appear to be antagonistic with the possibility of divergence by host-specialization within *M. oryzae*, our finding that the different lineages within *M. oryzae* tend to be sampled on a single host suggests that ecological barriers alone may in fact contribute to reduce gene flow substantially between host-specific lineages. Differences in the geographic distribution of hosts, for which the level of sympatry has varied -and still varies- in space and time, might also contribute to reduced gene flow between lineages infecting different hosts, although some level of sympatry at some time is required so that new hosts could become infected, triggering host-range expansion or host-shifting.

Mating within host (i.e. reproduction between individuals infecting the same host), and to a lesser extent mating system isolation (i.e. lack of outcrossing reproduction), may contribute to further reduce gene flow between *M. oryzae* lineages. The fact that mating in *M. oryzae* likely occurs within host tissues, such as dead stems [50], may participate in the maintenance of the different lineages by decreasing the rate of reproduction between isolates adapted to different hosts [6]. Loss of sexual fertility also appears to have a role in lineage maintenance. The rice lineage, in particular, is single mating-type and female-sterile throughout most of its range, which would reduce the chance of outcrossing sex with members of other lineages [51]. Our analyses rejected the null hypothesis of clonality in all lineages, but they provided no time frame for the detected recombination events. Population-level studies and experimental measurements of mating type ratios and female fertility are needed to assess the reproductive mode of the different lineages within *M. oryzae* in the field.

### Inter-lineage gene flow

Several potential barriers contribute to reduce genetic exchanges between *M. oryzae* lineages (see previous paragraph), but not completely so, as evidenced by signal of gene flow and admixture detected in our genomic data. We hypothesize that the lack of strict host specialization of the different lineages is a key driver of inter-lineage gene flow. Many of the grass or cereal species that are hosts to *M. oryzae* are widely cultivated as staple crops or widely distributed as pasture or weeds, including “universal suscepts” such as barley, Italian ryegrass, tall fescue and weeping lovegrass [40], increasing the chance for encounters and mating between isolates with overlapping host ranges. These shared hosts may act as a platform facilitating encounters and mating between fertile and compatible isolates from different lineages, thereby enabling inter-lineage gene flow [52]. Plant health vigilance is therefore warranted for disease emergence via recombination in regions where multiple lineages are in contact and shared hosts are present. This is particularly so, given that once infection of novel host has taken place (i.e. host shift or host range expansion), the fungus has the capacity to build inoculum levels very rapidly, facilitating spread of the disease over considerable distances. It is striking, for example, that wheat blast has, within a year, spread from Bangladesh into the West Bengal region of India where it emerged in 2017 (openwheatblast.org).

## Conclusion

Using a population genomics framework, we show that *M. oryzae* is subdivided into multiple lineages with limited host range and present evidence of genetic exchanges between them. Our findings provide greater understanding of the eco-evolutionary factors underlying the diversification of *M. oryzae* and highlight the practicality of genomic data for epidemiological surveillance of its different intraspecific lineages. Reappraisal of species boundaries within *M. oryzae* refuted the existence of a novel cryptic phylogenetic species named *P. graminis-tritici*, underlining that the use of node support in total evidence genealogies based on a limited dataset in terms of number of loci and of range of variation in origin (geography and host) of isolates can lead to erroneously identify fungal cryptic species. Our work illustrates the growing divide between taxonomy that ‘creates the language of biodiversity’ [53] based on limited sets of characters, and genomic data that reveals more finely the complexity and continuous nature of the lineage divergence process called speciation.

## Materials and Methods

### Fungal strains

Thirty-eight newly sequenced genomes were analyzed together with 43 published genomes [13, 14, 22, 54-56] resulting in a total of 81 *Magnaporthe* strains, including 76 *M. oryzae* genomes representing 12 different hosts available for analysis (Table 1). We also included as outgroups one strain of *Pyricularia pennisetigena* from *Pennisetum* sp. and four strains of *Pyricularia grisea* (syn. *Magnaporthe grisea*) from crabgrass (*Digitaria sanguinalis*). All newly sequenced strains were single-spored prior to DNA extraction.

### Genome sequencing and assembly

New genome data were produced by an international collaborative effort. Characteristics of genome assemblies are summarized in Table S3. For newly sequenced genomes provided by MF and BV, sequences were acquired on a MiSeq machine (Illumina, Inc.). Sequences were assembled using the paired-end mode in NEWBLER V2.9 (Roche Diagnostics, Indianapolis, IN). A custom perl script was used to merge the resulting scaffolds and contigs files in a non-redundant fashion to generate a final assembly. Newly sequenced genomes BR130 and WHTQ provided by TM were sequenced using an Illumina paired-end sequencing approach at >50X depth. Short reads were assembled *de-novo* using VELVET 1.2.10 [57] resulting in a 41.5Mb genome for BR130 with N50 44.8Kb, and 43.7Mb for WHTQ with N50 36.2Kb. For newly sequenced genomes provided by DS and NT, DNA was sequenced on the Illumina HiSeq 2500 producing 100 base paired-end reads, except in the case of VO107 which was sequenced on the Illumina Genome Analyzer II producing 36 base paired-end reads. Reads were filtered using FASTQ-MCF and assembled ’de novo’ using VELVET 1.2.10 [57].

Orthologous genes identification in genomic sequences

Protein-coding gene models were predicted using AUGUSTUS V3.0.3 [58]. Orthologous genes were identified in the 76-genomes *M. oryzae* or in the dataset including outgroups using PROTEINORTHO [59]. The v8 version of the 70-15 *M. oryzae* reference genome [60] was added at this step in order to validate the predicted sets of orthologs. Only orthologs that were single-copy in all genomes were included in subsequent analyses. Genes of each single-copy orthologs sets were aligned using MACSE [61]. Sequences from the lab strain 70-15 were removed and not included in further analyses due to previously shown hybrid origin [13]. Only alignments containing polymorphic sites within *M. oryzae* strains were kept for further analyses. This resulted in 2,241 alignments for the whole dataset, and 2,682 alignments for the 76 *M. oryzae* strains.

### Population subdivision, summary statistics of polymorphism and divergence

Population subdivision was analyzed using DAPC and STRUCTURE [30, 36-38], based on multilocus haplotype profiles identified from ortholog alignments using a custom Python script. DAPC was performed using the ADEGENET package in R [13]. We retained the first 30 principal components, and the first 4 discriminant functions. Ten independent STRUCTURE runs were carried out for each number of clusters K, with 100,000 MCMC iterations after a burn-in of 50,000 steps.

Polymorphism statistics were computed using EGGLIB 3.0.0b10 [62] excluding sites with >30% missing data. Divergence statistics were computed using a custom perl script. To infer total evidence trees within the 76 *M. oryzae* strains (respectively within the 81 *Magnaporthe* strains), all sequences from the 2,682 (respectively 2,241) orthologous sequences were concatened. The maximum-likelihood tree was inferred using RAxML [63] with the GTR-gamma model, and bootstrap supports were estimated after 1000 replicates.

### Retrieval of loci used in the Castroagudin et al. (2016) study

The 10 loci used by Castroagudin et al. [24], i.e. actin (ACT), beta-tubulin1 (βT-1), calmodulin (CAL), chitin synthase 1 (CHS1), translation elongation factor 1-alpha (EF1-α), hydrophobin (MPG1), nitrogen regulatory protein 1 (NUT1), and three anonymous markers (CH6, CH7-BAC7, CH7-BAC9), were search in all genomes using BLASTn. Due to heterogeneity in the quality of assemblies, 9 of the 10 loci could be full-length retrieved without ambiguity in 68 out of the 81 available genomes, still representative of the diversity of host plants.

### Secondary data analysis

Species recognition based on multiple gene genealogies as described by Castroagudin et al. [24] was repeated following the reported methods. The robustness of the Pgt species inference was tested by re-iterating the study, omitting one marker at a time. Individual genealogies were built using RAxML with the GTR-gamma model and 100 bootstrap replicates.

### Inference of “species tree” using ASTRAL

The ASTRAL method [27, 29] is based on the multi-species coalescent and allows taking into account possible discrepancies among individual gene genealogies to infer the “species tree”. Individual genealogies inferred using RAxML with the GTR-gamma model and 100 bootstrap replicates, were used as input data for ASTRAL analysis. Local supports around branches were evaluated with 100 multilocus bootstrapping using the bootstrap replicates inferred from each individual gene tree as input data, and with local quartet supports (q1, obtained using the −t option of ASTRAL) that represent the proportion of quartets recovered from the whole set of individual gene trees that agree with the local topology around the branch in the species tree.

### MPG1-based classification

The *MPG1* hydrophobin sequence is described as being diagnostic for the *P. graminis-tritici/M. oryzae* species split [24]. *MPG1* sequences from one of each species (GIs: KU952644.1 for *P. gramini-triticis*, KU952661.1 for *M. oryzae*) were used as BLAST [64] queries to classify isolates as either *P. graminis-tritici* or *M. oryzae*.

### Signatures of gene flow and/or incomplete lineage sorting

A phylogenetic network was built using SPLITSTREE 4.13 [65], based on the concatenation of sequences at single-copy orthologs identified in *M. oryzae*, excluding sites with missing data, sites with gaps, singletons, and monomorphic sites. The null hypothesis of no recombination was tested using the PHI test implemented in SPLITSTREE.

### ABBA/BABA tests

ABBA/BABA tests were performed using custom python scripts. The D statistic measuring the normalized difference in counts of ABBA and BABA sites was computed using equation (2) in ref. [66]. Significance was calculated using block jackknife approach (100 replicates, 1k SNPs blocks), to account for non-independence among sites.

### Probabilistic chromosome painting

We used CHROMOPAINTER program version 0.0.4 for probabilistic chromosome painting. This analysis was based on biallelic SNPs without missing data identified in the set of 2,682 single-copy orthologs, ordered according to their position in the reference genome of the rice-infecting strain 70-15. We initially estimated the recombination scaling constant N_e_ and emission probabilities (µ) by running the expectation-maximization algorithm with 200 iterations for each lineage and chromosome. Estimates of N_e_ and µ were then computed as averages across lineages, weighted by chromosome length, rounded to the nearest thousand for N_e_ (N_e_=5000; µ=0.0009). The file *recom_rate_infile* detailing the recombination rate between SNPs was built using the INTERVAL program in LDHAT version 2.2 [67] based on the whole dataset combining isolates from all lineages, with 10 repeats by chromosome to check for convergence. Estimated N_e_ and µ values and the per-chromosome recombination maps estimated using LDHAT were then used to paint the chromosomes of each lineage, considering the remaining lineages as donors, using 200 expectation-maximization iterations. For each lineage and each chromosome, CHROMOPAINTER was run thrice to check for convergence.

### Phylogenetic species recognition

We used an implementation of the GCPSR scalable to genomic data (<u>https://github.com/b-brankovics/GCPSR</u>). The method works in two steps: (1) Concordance and non-discordance analysis produces a genealogy that has clades that are both concordant and non-discordant across single gene genealogies, with support value for each of the clades being the number of single gene genealogies harboring the given clade at bootstrap support above 95%; (2) Exhaustive subdivision places all the strains into the least inclusive clades, by removing clades that would specify a species within potential phylogenetic species. We kept only two outgroup sequences per gene (BR29, *M. grisea;* Pm1, *M. pennisetigena*) to make sure to have the same isolate at the root of all genealogies (Pm1 isolate). Majority-rule consensus trees were produced from 100 outgrouped RAxML bootstrap replicates for all 2241 genes. The concordance and non-discordance analysis was carried out assuming 95 as the minimum bootstrap support value, and a discordance threshold of 1. Exhaustive subdivision was carried out using a concordance threshold of 1121.

### Whole genome alignment & tree building

A custom perl script was used to mask sequences that occur in multiple alignments when the genome is BLASTed against itself. The masked genomes were then aligned in a pairwise fashion against all other genomes using BLAST [64]. Regions that did not uniquely align in each pair at a threshold of 1e^−200^, were excluded. SNPs were then identified for each pairwise comparison and scaled by the total number of nucleotides aligned after excluding repetitive and duplicate regions. This produced a distance metric of SNPs/Mb of uniquely aligned DNA. The pairwise distances were used to construct phylogenetic trees with the neighbor-joining method as implemented in the R package, Analyses of Phylogenetics and Evolution (APE) [68].

Because alignments are in pairwise sets as opposed to a single orthologous set, assessment of confidence values by traditional bootstrapping by resampling with replacement is not possible. Instead, confidence values were assigned by creating 1,000 bootstrap trees with noise added from a normal distribution with a mean of zero, and the standard deviation derived from the pairwise distances between or within groups.

## Acknowledgments

We thank Sophien Kamoun for inspiration and for providing critical input on a previous version of the manuscript, Alfredo Urshimura for collecting and supplying us with the DNA of Brazilian isolates, and the Southgreen and Migale computing facilities. We also thank A. Akhunova at the Kansas State University Integrated Genomics Facility, and J. Webb, M. Heist and R. Ellsworth at the University of Kentucky for their technical assistance. Support is acknowledged by the Agriculture and Food Research Initiative Competitive Grant No. 2013-68004-20378 from the USDA National Institute of Food and Agriculture. This is contribution number 18-005-J from the Kansas Agricultural Experiment Station and contribution number XX-XX-XXX from the Kentucky Agricultural Experiment Station.

## Supplementary Tables legends

**Table S1. Pairwise distances measured in SNPs per megabase of uniquely aligned DNA**

**Table S2. (A) Gene flow signatures from ABBA/BABA tests.** P1, P2, P3 refer to the three lineages used for the tests. The D statistic tests for an overrepresentation of ABBA versus BABA patterns. SE is the standard error. Z-score and P-value for the test of whether D differs significantly from zero, calculated using 1000 block jackknifes of 100 SNPs. Analyses were based on 354,848 biallelic SNPs identified in 2241 single-copy orthologous genes, with *M. grisea* as the outgroup. Brachiaria1: Bm88324. Brachiaria2: Bd8401. Eleusine1 and Eleusine2 are, respectively, the light orange and orange clusters in DAPC analysis. Boldface labels indicate pairs connected by gene flow, as indicated by the sign of D. Time period represents the time scale over which gene flow is measured, as described in Figure S6. **(B) Timing of gene flow.** For all pairs of lineages belonging to triplets that yielded D values significantly different from zero, the corresponding time period over which gene flow was measured (as defined in Figure S6) is indicated. Each pair belongs to multiple triplets, spanning different time periods, and the reported time period is therefore the consensus of the corresponding time scales. For instance, the pair (brachiaria2, eleusine1) was included in triplets measuring gene flow at both intermediate (time periods 1 and 2) and recent (time period 1) time scales, and the consensus time period is therefore “1+2”.

**Table S3. Summary statistics of genome assemblies used in this study.**

## Supplementary Figure legends

**Figures S1A-S1J Maximum likelihood tree based on *MPG1, ACT1, b-tubulin1, BAC6, CAL, CH7BAC7, CH7BAC9, CHS, Ef1a* and *NUT1* marker, respectively.** Trees are represented as unrooted cladograms. Dark branches represent branches with bootstrap support > 50 after 100 bootstrap replicates (corresponding support are indicated). Clades are labeled according to the convention used by Castroagudin et al. [24]. Green dots: representatives of Pgt (*Pyricularia graminis-tritici sp. nov.*). Red dots: representatives of Pot (*Pyricularia oryzae* pathotype Triticum) clade 1. Blue dots: representatives of Pot clade 2. Orange dots: representatives of Poo (*Pyricularia oryzae* pathotype Oryza).

**Figure S2. Species tree inference based on the dataset of Castroagudin et al. [24] using ASTRAL.** The tree is represented as an unrooted cladogram.

Multilocus bootstrap supports above 50 are indicates above branches. Dark branches represent branches with bootstrap support > 50 after 100 bootstrap replicates (corresponding support are indicated). Number in brackets are q1 local quartet supports (i.e. the proportion of quartet trees in individual genealogies that agree with the topology recovered by the ASTRAL analysis around the branch). Clades are labeled according to the convention used by Castroagudin et al. [24] as in S1-S10 Figures.

**Figure S3. Analyses of population subdivision using clustering algorithms. (A) Bayesian Information Criterion vs number of clusters assumed in DAPC analysis.** The Bayesian Information Criterion assesses the fit of models of population structure assuming different K values. **(B) DAPC analysis of population subdivision, assuming K=2 to K=15 clusters.** Each isolate is represented by a thick vertical line divided into K segments that represent the isolate’s estimated membership probabilities in the K clusters. The host of origin of samples is shown below the barplot. **(C) Log likelihood of data vs number of clusters assumed in STRUCTURE analysis.** Error bars are standard deviations of likelihood across STRUCTURE repeats. **(D) STRUCTURE analysis of population subdivision, assuming K=2 to K=15 clusters.** Each isolate is represented by a thick vertical line divided into K segments that represent the isolate’s estimated membership proportions in the K clusters (note that two to seven clusters are empty, i.e. represented by no isolates, for models with K>9). The host of origin of samples is shown below the barplot.

**Figure S4. Neighbor joining tree showing the genetic distance separating the *M. oryzae* strains from *M. grisea* and *M. pennisetigena.*** Distances are in SNPs/kb.

**Figure S5. Distribution of MPG1, BAC6, β-tubulin1, CAL, CH7BAC7, CH7BAC9, EF1α and NUT1 alleles among *M. oryzae* isolates as indicated by mapping onto the neighbor-joining tree built using whole genome SNP data.** Alleles were identified by using a reference marker sequence for each gene, to search all the genomes using BLAST. Sequence variants are noted above each tree using the BLAST backtrace operations (BTOP) format.

**Figure S6. Time periods for gene flow covered by different triplets of lineages in ABBA/BABA tests.** Heterogeneity in divergence time between members of ((P1,P2),P3) triplets allows to examine gene flow at three time scales [33]: (A) triplets including the most recently diverged lineages as P1 and P2 (i.e. the *Triticum* and *Lolium* lineages, the two *Eleusine* lineages, or the *Oryza* and *Setaria* lineages) carry information about gene flow across relatively recent times, (B) triplets including, as P1 and P2, two lineages from the same main group of lineages (i.e. *Eragrostis / Eleusine1 / Eleusine2 / Triticum / Lolium or Brachiaria2 / Setaria / Oryza*, excluding (P1,P2) pairs already used in (A)) carry information about gene flow across intermediate times, and (C) triplets including, as P1 and P2, two lineages from different groups of lineages (i.e. *Eragrostis / Eleusine1 / Eleusine2 / Triticum / Lolium and Brachiaria2 / Setaria / Oryza*) and *Stenotaphrum* or *Brachiaria1* as P3 carry information about gene flow across a relatively long time period. All three graphs correspond to hypothetical cases in which the D statistic that measures imbalance between ABBA and BABA types indicates gene flow between P2 and P3 (i.e. positive D values). In (A) and (B) multiple possible topologies are shown, as P1, P2 and P3 can either belong to the same group of lineages or to different groups of lineages.

## Supplementary Text legends

**Text S1. Probabilistic “chromosome painting” analyses.**

